# Immunomodulatory Effects of a Cell Processing Device: Insights from Single Cell RNA Sequencing and Gene Set Enrichment Analysis

**DOI:** 10.1101/2025.10.03.680363

**Authors:** Christopher J. Pino, James Odum, Taylor Medendorp, H. David Humes

## Abstract

The Selective Cytopheretic Device (SCD), marketed as QUELimmune^TM^, is an autologous leukocyte processing extracorporeal device which promotes immunomodulation of a systemic hyperinflammatory states. To date, interrogation of the mechanism of action (MoA) of this cell-directed therapy has provided insight into therapeutic impact on neutrophils. However, characterization of monocytes has proven to be more difficult due to their dynamic plasticity, requiring more sophisticated analysis methods. Single cell ribonucleic acid sequencing (scRNAseq) methodology using 10x Flex is reported to further elucidate MO MoA in SCD therapy. Analysis strategies of unsupervised clustering and interrogation of specific genes of interest based on previous mechanistic insights from cell surface marker analysis and bulk RNA sequencing were utilized. SCD-treated monocytes were demonstrated to have lower expression of several proinflammatory gene products including *TNF* and *IL-6* and enrichment in anti-inflammatory gene markers, suggesting that SCD monocyte processing results in the release of monocytes with a less inflammatory phenotype. Calcium-dependent gene pathways are also downregulated, consistent with the low iCa environment of SCD therapy. Furthermore, analysis points to the involvement of key transcription factors such as AP-1 and NFkB. Taken together with preclinical and clinical data, scRNAseq confirms several aspects of the suspected SCD MoA on monocytes to transform them to a lesser pro-inflammatory phenotype.

## Introduction

The innate immunologic system, primarily circulating neutrophils (NE) and monocytes (MO), are a first line of defense to combat infection and to initiate tissue repair and recovery upon insult (1). Systemic hyperinflammatory response may occur if this process becomes excessive and dysregulated. Cytokine storm is a term often used to describe this derangement, where proinflammatory cytokines tend to be augmented (2). In this state, the equilibrium of monocyte subsets is dysregulated, reducing the proportion of anti-inflammatory non-classical monocytes (CD14^-^CD16^+^) by promoting pro-inflammatory classical (CD14^+^CD16^-^) and intermediate (CD14^+^CD16^low^) MO phenotypes (3). These pro-inflammatory MO release cytokines such as interleukin (IL)-1β, IL-6, and tumor necrosis factor (TNF)-α which promotes tethering and infiltration of monocytes into tissues, as well as additional MO recruitment, generating a positive-feedback loop of inflammatory amplification (3, 4). After infiltrating tissues, MO can differentiate into macrophages. Generally, classical and intermediate MO preferentially differentiate into M1 macrophages, and non-classical MO preferentially differentiate into M2 macrophages (5). Under ideal conditions, inflammatory M1 macrophages would eventually undergo apoptosis, or polarize to M2 anti-inflammatory macrophages to control the inflammatory response (5, 6). Dysregulation of macrophage polarization in disease states can lead to secondary injury to host tissue (5, 6). Generally, therapies have targeted individual cytokines, such as TNF-α, IL-6, or IL-1α through the use of monoclonal antibodies or non-specific removal strategies, such as sorbent-based technologies, to reduce cytokine loads (7, 8). Robust and durable clinical outcomes have not been demonstrated with these approaches (7, 8).

Several complex clinical hyperinflammatory disorders, such as sepsis, and autoimmune diseases are caused by dysregulation of immune cells. Instead of treating the byproducts of this dysregulation, such as therapies aimed at neutralizing or filtering cytokine levels, these disorders may be better addressed seeking novel therapies to return dysregulated immune cells to homeostatic function. One such approach is utilizing a device called the selective cytopheretic device (SCD) within an extracorporeal blood treatment circuit. The SCD is composed of a biocompatible polysulfone hollow fiber membrane within polycarbonate cylindrical housing. The surface area ranges from 1.0-2.5m^2^ depending on the therapeutic dose. A low-shear stress blood flow path around the bundled fibers is generated with administration of the SCD via an extracorporeal continuous kidney replacement therapy (CKRT) circuit. This promotes binding of NE and MO, which are in an activated state as a consequence of a systemic inflammatory disease process (9, 10). Immunomodulation of the sequestered activated leukocytes (LE) within the SCD occurs when clinical protocols for regional citrate anticoagulation (RCA) (11, 12) of the blood passing through the SCD exposes LE to a low ionized calcium (iCa) environment (0.2-0.4mM), promoting LE adhesion to the SCD membrane before reintroduction into systemic circulation. The SCD has been found to reduce systemic inflammation and improve clinical outcomes in several disorders with acute and chronic hyperinflammation, including sepsis (13–15), AKI (14, 15), ARDS and cytokine storm associated with COVID-19 (10, 16), acute and chronic liver failure (9), cardiorenal syndrome (17), and others (18).

A previous research article published by our group explored SCD cell-directed therapy to phenotypically alter circulating LE of the innate immune system with a focus on the role of NE and MO using cell-surface marker characterizations and analysis of cytokine secretory rates (19). Activated NE adhere to SCD fibers and degranulate, releasing constituents of their exocytotic vesicles (19). Exposed to the low iCa environment, adhered neutrophils display characteristics of apoptotic senescence, which are then released and return to circulation to the bone marrow to diminish NE release (19). Due to the complexity of phenotypic classification of MO populations using cell-surface markers, as well as the relatively lower proportion of MO in the overall LE population, MO analysis was more limited. The results suggested that a pro-inflammatory subsets of circulating MO selectively bind to the SCD (19). Over time, a subset of the bound monocytes with a less inflammatory functional phenotype are then released from the membrane back into circulation. In this study, we aim to further expand upon these results using single-cell RNA sequencing. Single-cell RNA sequencing (scRNAseq) has been used by other groups to interrogate the therapeutic impact of extracorporeal circuits on circulating LE. Of specific relevance to this work, RCA has been shown to prevent the upregulation of surface activation markers, like human leukocyte antigen class II (HLA-DR) and produce alterations in expression of several MO gene families related to cell cycle regulation, cellular metabolism, and cytokine signaling (20). Because SCD therapy utilizes RCA to create a low iCa environment, the hypothesis that SCD therapy may elicit a similar response is logically consistent. In addition to calcium dependent gene pathways, expression changes of genes involved in critical regulatory pathways, such as those involving protein (AP)-1, nuclear factor kappa-light-chain-enhancer of activated B cells (NFκβ), and signal transducer and activator of transcription (STAT) proteins (5), may provide detailed insight into the specific immunomodulatory effects of SCD therapy and its potential downstream impacts on monocyte-derived macrophages (21). Proteins from all three of these families are involved in signaling pathways involved in inflammation, proliferation, differentiation, and cell-death (22). In order to interrogate how SCD therapy modulates gene expression of calcium-dependent and inflammatory genes and gene pathways in MO, scRNAseq was performed. The results support previous findings by our group that SCD released cells are of a less inflammatory phenotype and identify potential future focus areas for SCD MoA studies.

## Methods

### Human Blood collection

With our group’s experience conducting *in vitro* blood circuit (IVBC) studies (13, 19), it was understood that fresh human blood would be required to study the molecular underpinnings of LE activation and SCD MoA. Fresh animal blood has been found to be adequate for some cell surface marker evaluations (13), however, most ample sources of abattoir blood (porcine, bovine) are not amenable for scRNAseq analysis.

Banked blood or commercially sourced human blood which must be drawn, tested and shipped, with >24h delivery times were also ruled out for not being fresh enough (19). Due to these constraints, it was recognized that from a single human donor, the fresh blood draw volume would have to be limited, which would also necessitate limited circuit volumes. To limit therapy circuit volumes, a miniature version of the SCD and miniaturized circuits were utilized only requiring 50-100mL. Blood was collected directly into heparin (∼10U/mL). To achieve an iCa of 0.25-0.4mM, citrate in the form of ACD-A (Baxter, Deerfield IL) was titrated, generally requiring approximately 1:60 by volume (19). All protocols related to human blood collection from human donors were approved through the University of Michigan (UM) Institutional Review Board (IRB), HUM00209396. All human blood studies were carried out in accordance with the associated guidelines and regulations, and informed consent was obtained from all blood donors.

### Construction of miniature SCD for use during *in vitro* human blood circuit (IVBC) studies

Construction of miniature SCD with an approximate fiber surface area of 0.1m^2^ and a fill volume of approximately 20mL were constructed using as similar materials as possible to clinical devices. Please refer to Westover et al 2024 for details of miniature SCD construction (19).

### *In Vitro* Blood Circuit (IVBC) with Plasma Chase with Human Blood

Schematically, these studies are concisely described in Figure 1A. Blood is perfused through the devices allowing LE to adhere to the fibers within devices. Devices are rinsed to remove residual blood. Plasma is perfused through the device to determine the number and types of cells that detach over time. At the end of perfusion, any remaining cells are removed from devices to using a dissociation buffer. A similar IVBC model system was previously utilized in Westover et al (19). Please see Westover et al for further experimental details of circuit preparation, blood recirculation, plasma chase, and SCD elution. In the model system presented herein, cell isolated from blood were further enriched for MO, and enriched MO/NE populations from SCD were fixed and cryopreserved for scRNAseq via 10x Flex.

**Figure 1.**
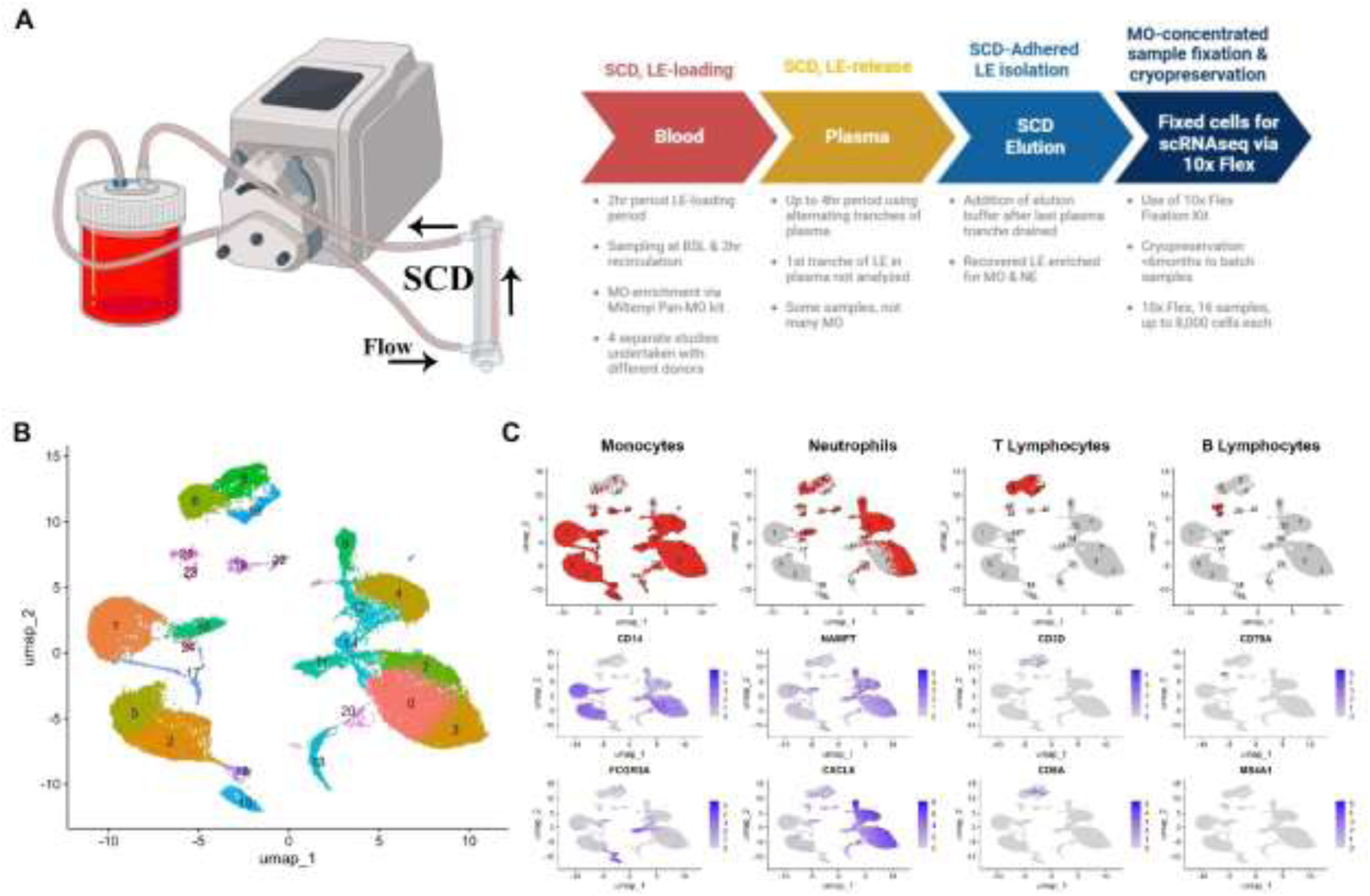
In vitro experimental design using miniaturized Selective Cytopheretic Device (SCD). **(A)** Schematic depiction of integrated single cell RNA sequencing dataset generated from human donor blood across 4 sample types: baseline, 2 hours of circulation through SCD (Circ), 4-hour chase with human plasma (Plasma), and elution (Elute). **(B)** UMAP representing 25 subpopulations of immune cells generated from in vitro SCD exposure integrated across all four sample types. **(C)** singleR (red) cell type assignments for monocytes, neutrophils, T lymphocytes, and B lymphocytes compared to expression of canonical genes (purple) for those respective cell types.

Circuit preparation: In brief, donor blood was transferred to 1L EVA bag (Baxter) with inlet and outlet access ports to enable blood flow. After adding citrate in the form of Anticoagulant Citrate Dextrose-Solution A (ACD-A, Baxter) to reach the prescribed low iCa range (0.25-0.4mM, targeting 0.3mM) for testing, blood was warmed to 37°C within a Hera cell culture incubator with CO2 injection disabled. Blood bags were gently mixed by hand throughout study durations to prevent plasma separation. Recirculating blood circuits were established by priming the miniature circuit and miniature SCD with normal saline (0.9% NaCl) to wet the circuit, followed by emptying the circuit, so as to not appreciably dilute the low volume of blood, once the blood bag was connected. Any residual normal saline (0.9% NaCl) or very dilute blood was pumped single pass to waste, and recirculation of blood was established thereafter.

Blood Recirculation Period: In brief, blood was recirculated for 2 hours with a MasterFlex model 7523-80 roller pump with size 17 BPT tubing for the pump segment, using a blood flow rate (BFR) of 10mL/min. The rest of the miniaturized circuit was comprised of silicone tubing with an inner diameter (ID) of 2.79mm (Cole-Parmer). This BFR in the miniature SCD mimics shear stresses in full-size clinical devices which have greater fiber surface areas as well as higher fill volumes. Blood samples were taken from well-mixed blood at baseline, as well from blood after 2 hours of recirculation. Blood samples were placed directly in 1.3mL ETDA vials (Sarstedt) and placed on ice for simultaneous batch processing later on with other samples. At the end of the recirculation period, the SCD and miniaturized SCD circuit were rinsed free of blood with normal saline and drained.

Plasma Chase Period: In brief, pooled human plasma generated by apheresis from heparinized blood was procured from Innovative Research (Novi, MI) and stored at -80°C up until use. Plasma stock was thawed, and ∼160-200mL of plasma was ultra-centrifuged to remove lipid aggregates. Similar to blood, citrate in the form of ACD-A was used to decrease iCa into the target range (0.25 – 0.4mM, targeting 0.3mM), mixing well and splitting the plasma into to two separate tranches of 80-100mL each. The first plasma chase period (0-60 minutes) utilized plasma tranche 1 . After recirculating through the SCD for 1h, at a rate of10mL/min (Fig 1A), this plasma stock was entirely removed from the SCD circuit, split into two 50mL conical tubes, and centrifuged at 500 x g to collect SCD-released LE. Plasma tranche 1 supernatant was removed and saved to be utilized later as required, leaving only a cell pellet for later analysis. Cell pellets were kept on ice, for simultaneous batch processing at the end of the study. Plasma tranche 2 was used in the second plasma chase period (60-120 minutes) and was processed with the same methodology as tranche 1: centrifuged, saving the plasma supernatant, and collection of the cell pellet. Plasma tranche 1 was reused in the third plasma chase period (120-180 minutes), and plasma tranche 2 was reused in the fourth plasma chase period (180-240 minutes), completing a total plasma chase of 4 hours as required. After the last period of plasma recirculation, and the last tranche of plasma collected, the SCD was rinsed with normal saline prior to SCD elution.

SCD Elution (Cell Dissociation) and Processing: To remove the adhered leukocytes from the miniature SCD, Normal saline was drained and replaced with 0.2% EDTA in PBS at a pH of 7.2 which causes LE to dissociate from SCD fibers. The miniature SCD was capped and incubated at room temperature for 30 minutes. Cell detachment was encouraged with vigorous shaking and gentle tapping of the device. Syringes were used to collect the spent elution buffer filled with detached LE from SCD (referred to as SCD elution).

Repetitive rinse steps with injecting fresh elution buffer and removing spent elution buffer were employed to remove remaining LE. Previous analysis of this methodology has demonstrated that few residual cells are not removed by elution process. Total elution volume was noted and split into conical tubes for centrifugation (500 x g, 10 minutes) to isolate cell pellets, which were combined.

In previous studies in the same model system, the total number of cells adhered on SCD were assessed by fluorescent labeling of nucleated cells with Hoerscht 33342 and counted while viewed with a hemacytometer. Differential counts of NE and MO were determined by Cytospin analysis using cytospin funnel In this work, due to relatively low numbers of cells available in plasma chase and SCD elution samples, this additional analysis was not enabled for all studies.

### Monocyte isolation

MO were isolated from baseline (BSL) and 2h circulation (CIRC) whole blood samples through a multistep process. First, red blood cells (RBC) were lysed using ammonium chloride lysis, at a ratio of 1mL per 9mL of lysis buffer. Isolated LE were then further enriched for MO using the Miltenyi Pan-MO kit following the kit’s instructions for magnetic separation using the MACS system. MO enrichment was assessed utilizing cytospin analysis using a cytospin funnel.

Red blood cell (RBC) contamination in SCD elution and plasma chase cell pellets was removed by ammonium chloride lysis, at a ratio of 1mL per 9mL of lysis buffer. After rinse steps, and resuspension in buffer a small volume of samples were taken to estimate total cells available by trypan blue counts aided by a hemocytometer. All cell pellets for plasma chase and SCD elution were enumerated by trypan blue counts.

### Cell fixation and storage

Instructions were followed associated with utilizing the 10x Flex Fixed Cell scRNAseq kit. Samples were stored at -80°C in Ultra Low Freezers (ULF) for no longer than 6 months per 10x instructions.

### Single cell RNA sequencing (scRNAseq)

Frozen cell samples were transported to the University of Michigan Advanced Genomics Core (AGC) for scRNAseq preparation and analysis. A cell number of 8,000 cells per sample, from sixteen total samples was targeted with a minimum number of cell reads set at 10,000. Gene count matrices were generated from raw fastq files via Cell Ranger (v8.0.1).

### scRNAseq analysis

Gene count matrices were loaded into R (4.2.1) using the Read10X function contained in Seurat (5.1.0, https://github.com/satijalab/seurat). A single Seurat object was generated from a total of 16 samples composed of 4 baseline (BSL), 4 circulation (Circ), 4 plasma chase (Chase), and 4 eluted (Elute) samples. Standard quality control ensured only cells with 200-6500 features and <30% mitochondrial RNA were included for analysis. Data was subsequently scaled by a factor of 10,000 and Log10 normalized. FindVariableFeatures was employed to identify 2000 highly variable genes. A linear transformation was performed prior to the principal component analysis. Batch correction to individual samples was performed using Harmony (1.2.1). Next, nearest neighbors was identified using FindNeighbors with 20 principal components selected as appropriately representative of the dataset. The cells were subsequently clustered using FindClusters. Cell type identities were assigned using SingleR (1.10.0) with reference to HumanPrimaryCellAtlasData (23), and monocytes were subsetted for targeted analysis. The subsetted monocytes underwent unsupervised clustering after repeat scaling, principal component analysis (20 dimensions), and Harmony (24) batch correction with a resolution of 0.6. FindAllMarkers was employed to identify the top genes associated with each MO cluster.

### Classical and Non-Classical Monocyte Scoring

Classical and Non-Classical Monocyte Scores were generated using the AddModuleScore function in Seurat based on a list of top 10 genes closely associated with each of these monocyte phenotypes (Supplemental Table). Briefly, AddModuleScore (25) calculates the average expression of each prespecified gene set across all cells in the cluster weighted to the aggregate expression of randomly selected control features. These scores were graphically represented using violin plots.

### Pathway Analysis

For Gene Ontology (GO) Enrichment Analysis, GO: Biological Process pathways were selected that aligned to key proinflammatory, anti-inflammatory, calcium handling, and transcription factor mediated pathways. ClusterProfiler (26) was employed to perform analysis for curated GO: Biological Process pathways across subsetted MO clusters. Heatmaps were generated to depict the relative expression for each GO pathway per MO cluster with unsupervised clustering.

### Slingshot Pseudotime Analysis

A pseudotime analysis was performed on the subsetted MO clusters utilizing Slingshot (27) based on pre-computed harmony embeddings with unsupervised identification of the starting cluster performed by Slingshot. Next, edgeR (28) was employed to generate the differential gene expression across MO clusters according to pseudotime values, and ggplot2 was subsequently used to plot relative gene expression of targeted genes across pseudotime values.

### Interrogation of Calcium Signaling Related Gene Expression in MO clusters

Utilizing the clusters established for only the MO subpopulation identified by the process detailed above, several genes of interest were specifically interrogated for their involvement with calcium related processes which may be impacted by the low ionized calcium environment of SCD therapy. Genes to interrogate included ORAI1, ORAI2, ORAI3, STIM1 and STIM2, members of a calcium signaling pathway called store-operated calcium entry (SOCE) (29). Calcium-release activated calcium (CRAC) channels are mediated by the ORAI/STIM complex. In this complex, ORAI proteins are pore-forming proteins, with the calcium store sensor being the stromal interaction molecules (STIM isoforms) (29). Plasma membrane calcium ATPase (PMCA) pump isoforms ATP2B1 and ATP2B2 and the sodium/calcium ion exchanger (NCX) are involved in on the plasma membrane to pump calcium out of cells, to provide a low cytosolic concentration allowing for calcium mediated signaling (30, 31). Mitochondrial calcium uptake protein isoforms MICU1 and MICU2 were analyzed. Lastly, NADPH oxidase 2 (NOX-2) was interrogated which can mediate oxidative burst in monocytes (32).

### Interrogation of expression for genes of interest in MO clusters for targets previously identified by bulk RNAseq

In previous studies, bulk RNAseq was utilized as a preliminary evaluation for transcriptional changes. In those very limited studies simply looking at baseline blood LE and those treated with a citrate to create a low iCa environment, a downregulation of the CCL subfamily (CCL2,7,8,12, 15 and 23) and CCR7 was observed, as well as the CXCL family (CXCL9 (Mig), CXCL10 (IP-10), and CXCL11 (I-TAC) and IL12 family cytokines (IL12, 27, 35) (previously unpublished data, data not shown). An increase was seen in MCP-1 (previously unpublished data, data not shown). These targets were specifically interrogated in scRNAseq, to see if there was confirmation of expression shifts in this IVBC model system.

### Interrogation of Apoptosis and Senescence Related Gene Expression in MO clusters

In studying SCD mechanism of action in NE, it was found that SCD-processing of cells to become pro-apoptotic or senescent was a major change identified (19). Therefore, related targets were specifically interrogated in scRNAseq, to see if there was confirmation of expression shifts in this IVBC model system.

### Unsupervised MO Pairwise Comparisons by *In Vitro* Stage

DESeq2 (1.36.0) was used to perform multiple pairwise comparisons across differentially expressed genes among subsetted MO according to the *in vitro* stage of cell isolation. Volcano Plots were generated with thresholds p < 0.01 and log2Fold change > 0.5. ToppGene was utilized to query differentially regulated Gene Ontology (GO):Biological Process pathways based on the list of differentially expressed genes in each pairwise comparison (33). Representative GO pathways were selected to reduce redundancy and highlight pathways related to inflammation, infection, wound healing, cell cycle, metabolic reprogramming, autophagy, and/or apoptosis.

## Results

The data reported herein is meant to be interpreted from the vantage point of several previous pre- clinical and clinical investigations, which have demonstrated the selectivity of the adhesion of activated, circulating LE on the SCD fibers membranes. The studies have shown that NE and MO, but not lymphocytes, are the dominant cell types that adhere to these membrane materials (9, 15, 16). Furthermore, predominantly pro-inflammatory MO (generally CD11b+, CD14+) that tend to be classical monocytes (CD14+ CD16low) or intermediate (CD14+ CD16+) adhere from the circulation during a hyper-inflammatory state (9, 16). Non-classical MO (CD14low CD16+) are anti-inflammatory (34, 35), which are also known as patrolling MO. In the IVBC model system used in this current work, changes were not observed in the overall distribution of monocyte subtypes when comparing baseline (BSL) to the 2hr recirculation (CIRC) blood samples (19).

However, the percentage of non-classical monocytes in the BSL (6.0%) and CIRC (5.6%) were significantly higher than in the bound cells (ranging from 0.7-1.4%, p<.05) indicating that few non-classical monocytes adhere to SCD fibers (19).

### Unsupervised Clustering Results

The results of unsupervised clustering of all cells are shown in Figure 1B, with a total of 24 clusters. Given significant overlap between gene signatures of MO and other immune cells in several clusters, singleR was employed to assist in cell type identification of each cell in the dataset. Presumed MO, NE, T-lymphocytes and B-lymphocytes as identified by single R were compared with canonical markers for each (Figure 1C). Overlap of these canonical markers confirms potential intermixing of NE and MO within some of these clusters, suggesting strong similarities in cell expression profiles in cases.

Results of unsupervised clustering of the 25877 cells assigned as MO by singleR with reference to HumanPrimaryCellAtlasData generated 11 MO clusters (Figure 2A, Supplemental Figure 1). Panel 2B shows the contribution of cells in that UMAP space based on the experimental phase of the cell samples: BSL, Circ, Plasma Chase and SCD Elution. BSL cells correlate most with clusters 2, 3, 4, 7, and 10 on the right side of the UMAP, while Circ samples correlate most with clusters 0, 1, 8, and 9 on the left side of the UMAP (Figure 2B, Figure 4B). Due to the relatively low number of cells analyzed from Plasma Chase and Elute samples (Supplemental Figure 1), these cells are found in smaller proportions in their associated clusters, but are present most in clusters 0, 1, 6, 8, 9, and 11 (Figure 2B, Figure 4B). Clusters 5 and 6 have a more mixed composition of cells from experimental phases. Cluster 5 contains a mix of cells from BSL, Circ, and a small proportion of Plasma Chase cells, while Cluster 6 has a mix of cells from all four experimental stages (Figure 4B). The top 5 marker genes per monocyte cluster are shown in a heatmap in panel 2C. Notably, chemokines and chemokine receptors, such as *CXCL1*, *CCR5*, *CCL4L2*, and *CCL22*, are enriched in some clusters, as well as *FOS,* a subunit of the AP-1 transcription factor complex). Looking at the UMAP space using cell markers *CD14* and *CD16* (*FCGR3A*) suggests that cells in cluster 5 are low in *CD14* and positive for *CD16*, consistent with a non-classical MO phenotype (Figure 2D). This is further evidenced by classical and non-classical MO scores, where cluster 5 has the lowest classical MO score and the highest non-classical MO score (Figure 2E). This pattern is true but to a lesser extent with cluster 8, suggesting cells in this cluster may also have a non-classical phenotype (Figure 2E). The clusters with the highest classical MO scores appear to be BSL associated clusters 2, 3, 4, and 7, which supports a reduction in monocytes with a classical phenotype following SCD treatment.

**Figure 2.**
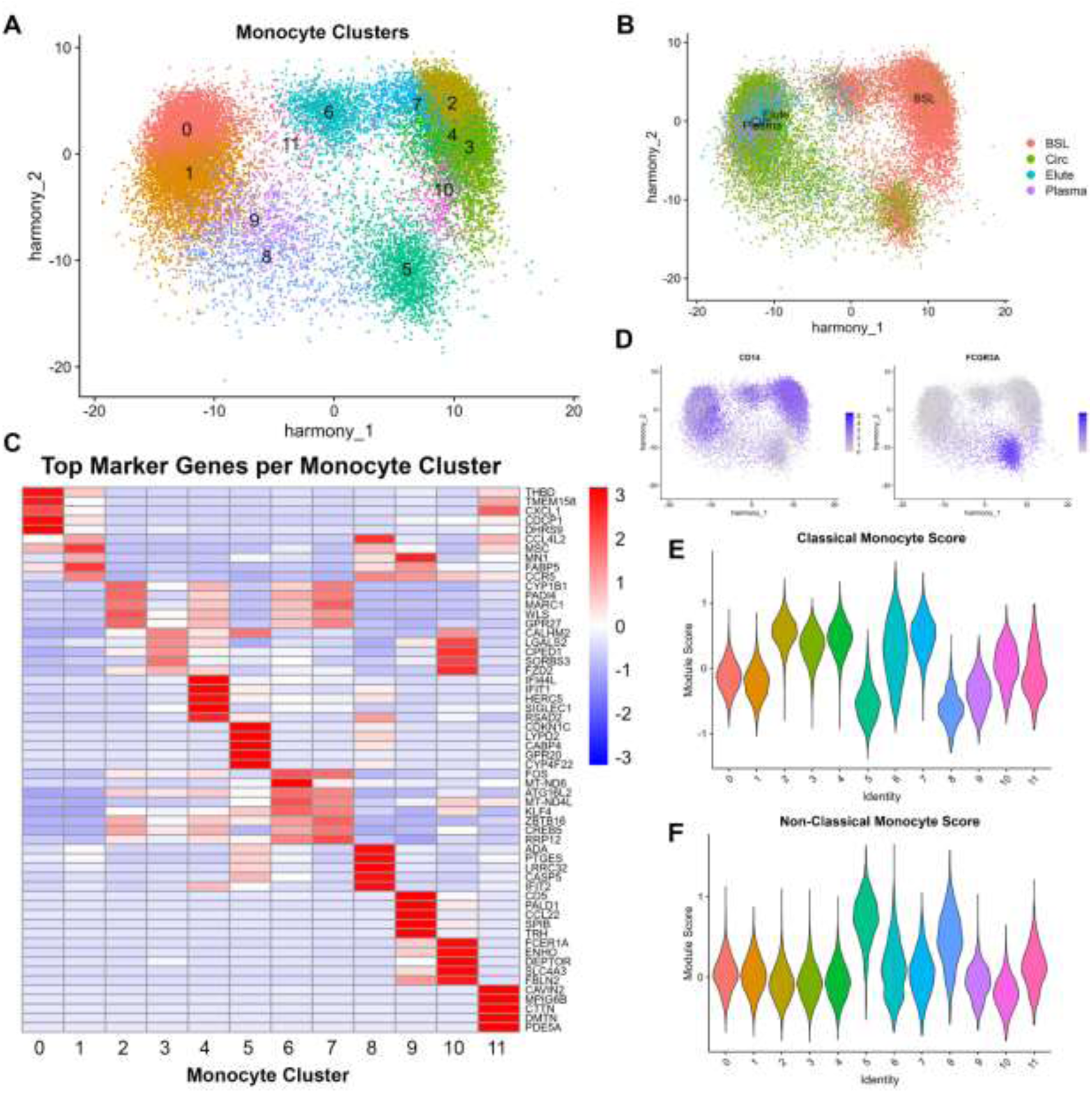
Monocyte subpopulations revealed during exposure to SCD. **(A)** UMAP representing 12 monocyte subpopulations across the integrated dataset consisting of BSL, Circ, Plasma, and Elute stages, batch corrected using Harmony reduction. **(B)** UMAP depicting the contribution of each stage across monocyte subpopulations. **(C)** Top 5 genes expressed across each monocyte cluster. **(D)** Expression of *CD14* and *FCGR3A* (CD16) across monocyte subpopulations reveals candidate clusters for Classical (CD14^+^/CD16^-^) and Non-Classical (CD14^-^/CD16^+^) monocytes. **(E)** Classical Monocyte Scores were generated for each monocyte cluster via AddModuleScore in the Seurat package for top 10 genes closely associated with Classical Monocytes (25). **(F**) Non-Classical Monocyte Scores were generated for each monocyte cluster via AddModuleScore in the Seurat package for top 10 genes closely associated with Non-Classical Monocytes.

Analysis of pro- and anti-inflammatory genes and pathways by cluster are summarized in Figure 3. Pro-inflammatory cytokines *IL-6* and *TNF-α* are most expressed in cells from clusters 8 and 11, followed by clusters 0 and 1 which appear to have much lower, but non-zero expression (Figure 3A). Anti-inflammatory genes *TGFB1* and *IL1RN* are expressed at varying levels in most MO clusters, with the highest expression levels occurring in clusters 0, 1, 9, and 11. There is a notable lack of expression of both *TGFB1* and *IL1RN* in some clusters (6 for *TGFB1,* 4, 5, and 6 for *IL1RN).* GO pathways, both pro-inflammatory and anti-inflammatory demonstrate marked differences in MO clusters, suggesting that the model system does provide activation via the mechanical pump utilized and substantial processing effects due to SCD treatment (Figure 3C, 3D).

**Figure 3.**
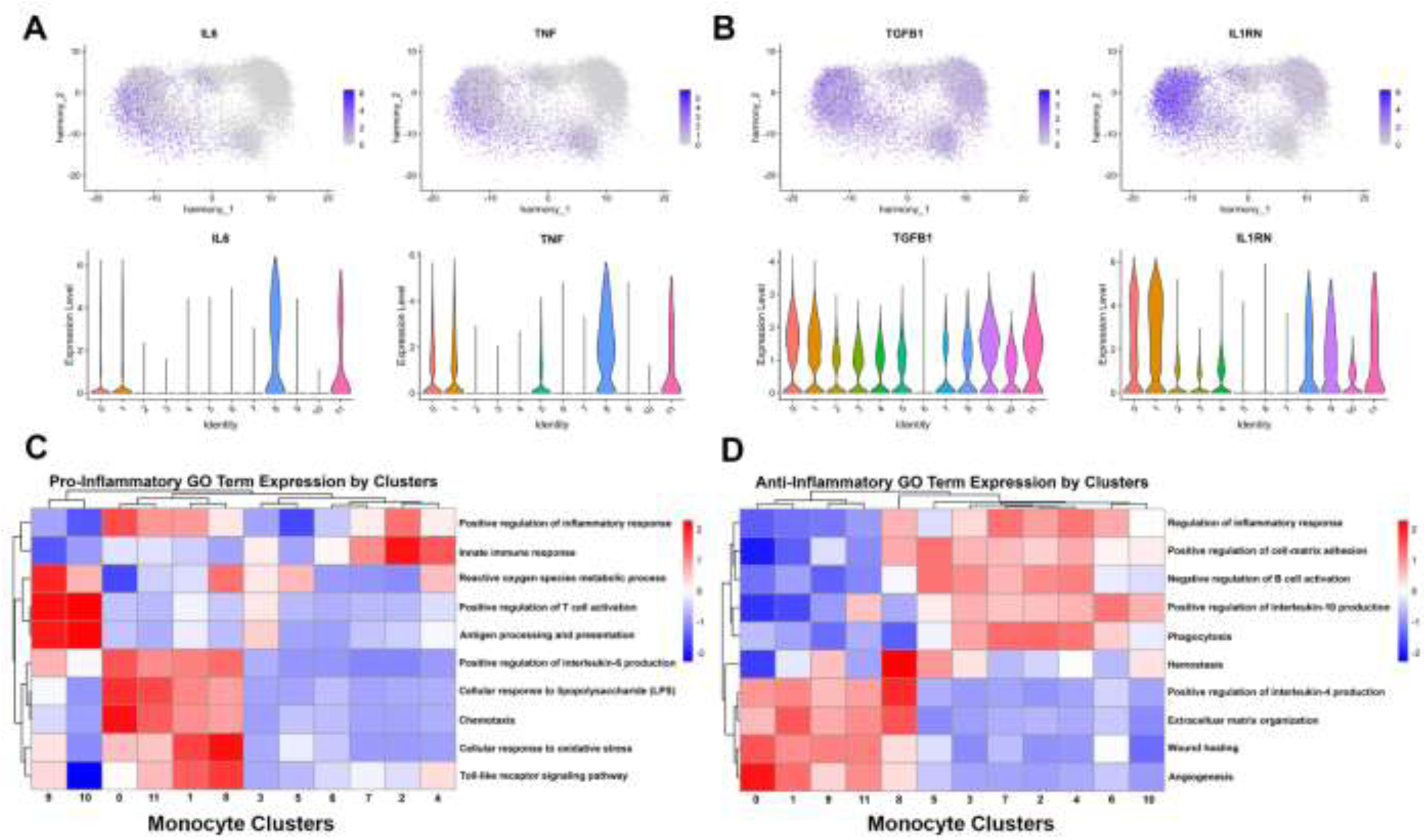
Inflammatory signatures across monocyte subpopulations. **(A)** Proinflammatory gene expression of *IL6* and *TNF* depicted in Feature Plots (purple) across monocyte subpopulations. Individual monocyte cluster expression of *IL6* and *TNF* are highest in clusters 0, 1, 8, and 11 as depicted in Violin Plots. **(B)** Anti-inflammatory gene expression of *TGFB1* and *IL1RN* depicted in Feature Plots (purple) across monocyte subpopulations. Individual monocyte cluster expression of *TGFB1* and *IL1RN* are increased across multiple monocyte clusters. **(C)** Heatmap representing Gene Ontology: Biological Process pathways across monocyte clusters. Pathways were selected for their associations with proinflammatory mechanisms. **(D)** Heatmap representing Gene Ontology: Biological Process pathways across monocyte clusters. Pathways were selected for their associations with anti-inflammatory mechanisms and wound healing.

**Figure 4.**
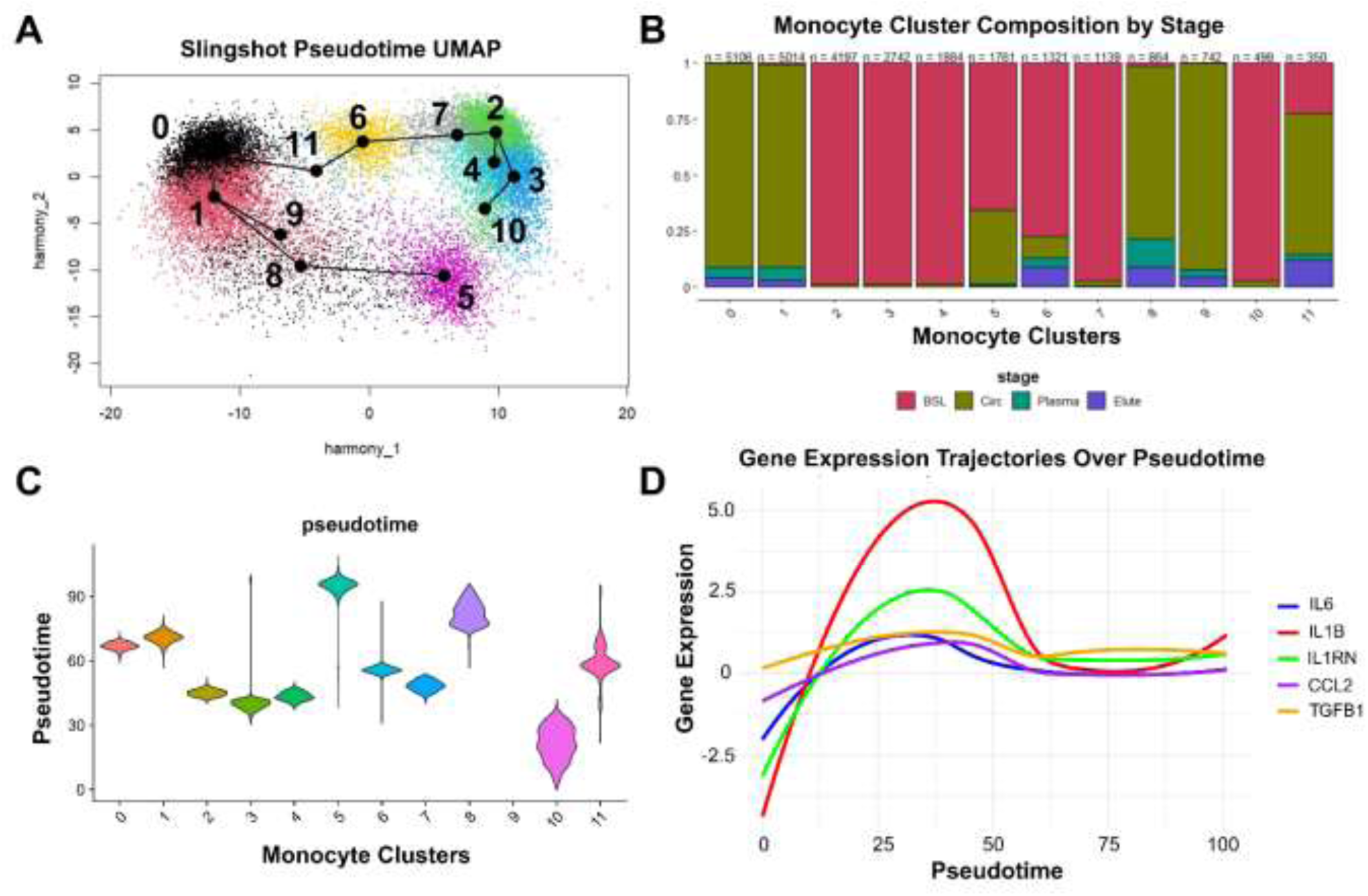
Identification of Monocyte Transcriptional Plasticity and Expression of Inflammatory Genes as a Function of Pseudotime. **(A)** Slingshot pseudotime analysis shows the computationally derived progression of monocytes over experiment duration super-imposed on UMAP with monocyte cluster numbers. Cluster 10 identified as the starting cluster during unbiased pseudotime analysis. **(B)**. Monocyte cluster composition by experimental stage (BSL, Circ, Plasma, Elute). **(C)** Violin plot depicting pseudotime values by monocyte cluster. **(D)** Dynamic changes in gene expression of targeted pro- and anti-inflammatory genes across aggregated monocytes over pseudotime.

Pseudotime analysis was performed, which utilizes known developmental and maturation markers to track suspected progression through clusters. Pseudotime initiated using BSL samples as a starting point progress through central clusters to far-left clusters, mirroring progression through the in vitro SCD modeling experiment, with MO activation stimulated by exposure to mechanical shear stress induced by the pump and/or interactions with the plastic interface of the tubing. Then, there is a return towards the right side of the UMAP space, with cluster 5 being the final “timepoint” (Figure 4A, 4C). This analysis suggests that a subpopulation of Circ cells that have been processed by SCD are altered to or remain associated with a non-classical MO phenotype (cluster 5) similar to what are found in the baseline population (Figure 4A, 4B). Because pseudotime appears to mirror progression of MO through the model system, analysis of trajectories of gene expression over pseudotime was performed using selected inflammatory related genes *IL-6, IL-1B, IL-1RN, CCL2, and TGFB1*. Gene expressions for *IL-6, IL-1B, IL-1RN,* and *CCL2* starts with an increase as pseudotime progresses from the right side of the UMAP to the left, followed by a decrease and plateau (Figure 3D). *IL-1B* shows the most dramatic modulation over pseudotime, followed by *IL-1RN*. Gene expression of *TGFB1* remains relatively steady throughout the progression of pseudotime.

Pro- and anti-inflammatory genes and pathways are further explored in Figure 5, now specifically utilizing BSL, Circ, Plasma Chase and Elute sample groupings (aggregated by pseudo-bulking individual cells by phase of sample collection) rather than the unsupervised MO clusters. Expression of pro-inflammatory cytokines *TNF, IL-1A, IL-1B, IL-6,* and *CXCL8* is lowest in BSL cells relative to other sample groups (Figure 5A). Both *TNF-α* and *IL-6* show relatively higher expression in Elute cells relative to BSL and Circ cells. *CXCL9*, *CXCL10*, and *CXCL11* show a reverse pattern, with expression highest in BSL relative to other experimental phases. Very little change in expression of these cytokines is seen between the other three sample groups. Interestingly, *IL12B* is strongly enriched in the Elute cell population relative to other sample populations, while the gene encoding one of its binding partners *IL23A* is strongly downregulated relative to other sample populations. Anti-inflammatory cytokine *IL-10* is most enriched in cells from Plasma Chase and Circ cells. Analysis of inflammatory GO pathways by experimental phase show intermixing of pro- and anti-inflammatory pathways (Figure 5B). The heatmap shows stark changes in pump activated and SCD-processed cells from baseline cells. Elute cells are enriched in pro-inflammatory pathways such as chemotaxis, IL-4 production, and toll-like receptor signaling pathways relative to all other groups but also show enrichment in anti-inflammatory pathways such as angiogenesis and wound healing (Figure 4B). This suggests that immunomodulation of SCD therapy is not completely immunosuppressive.

**Figure 5.**
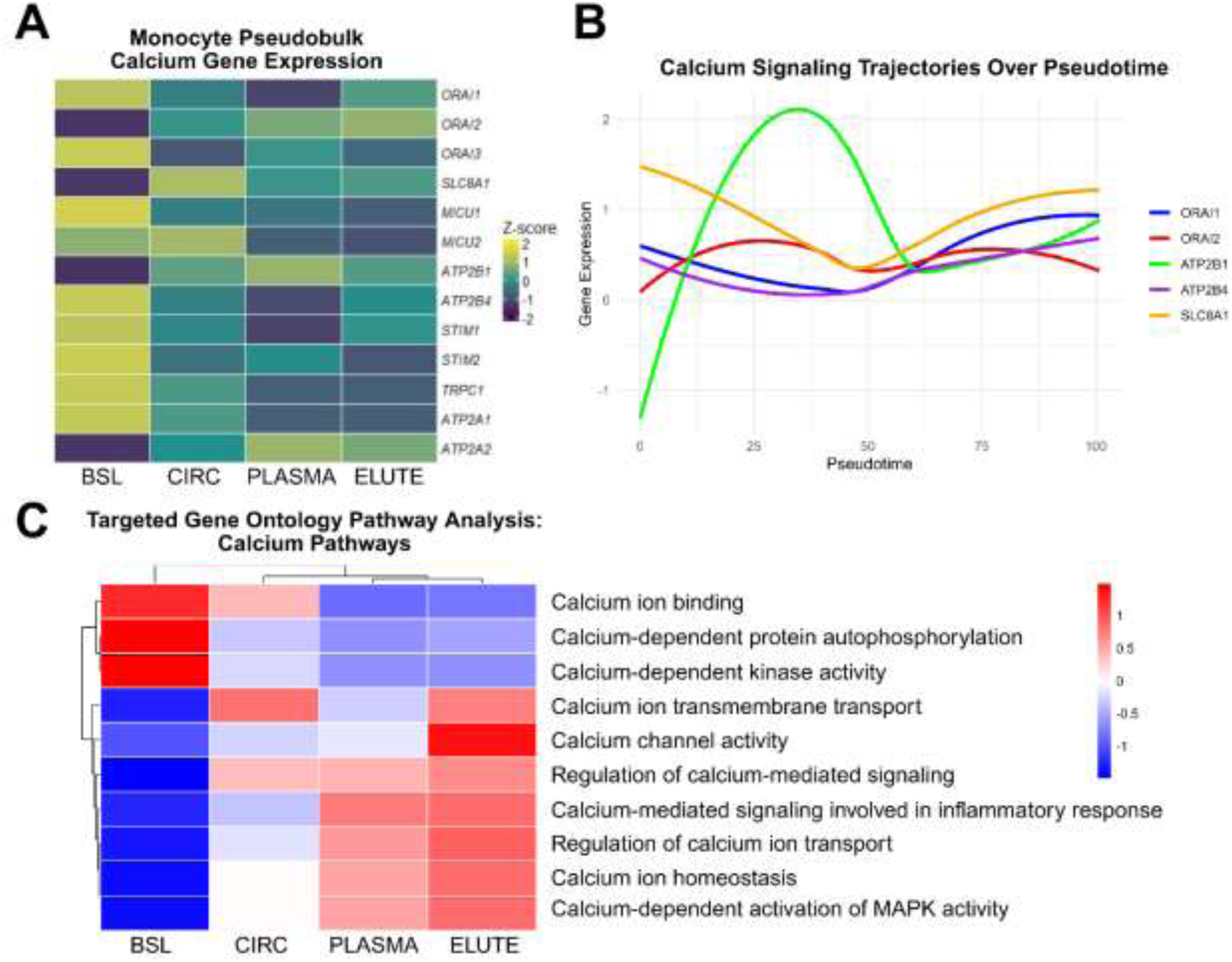
Dynamic changes in calcium related gene expression in monocytes by stage. **(A)** Heatmap depicting expression of targeted calcium-related genes in pseudobulk of monocytes by stage. **(B)** Expression of individual calcium related genes plotted across pseudotime indicates *ATP2B1* and *ORAI2* initially increase in unstimulated human blood at the onset of experimental exposure to SCD whereas *ORAI1*, *ATP2B4*, and *SLC8A1* initially decrease. However, all genes converge to follow a similar trajectory in the latter half of pseudotime. **(C)** Heatmap representing Gene Ontology: Biological Process pathways across BSL, Circ, Plasma, and Elute. Pathways were selected for their associations with calcium transport, calcium handling, and calcium mediated signaling.

### Results from Interrogation of Calcium Signaling Related Genes

Figure 6 aims to examine how SCD therapy and its low iCa environment impacts gene expression of calcium-dependent channels and signaling pathways. Genes of the ORAI and STIM families were assessed due to their role in the store-operated calcium entry (SOCE) mechanism, which resupplies store organelles (such as the endoplasmic reticulum) with Ca^2+^ after they are depleted (20). Additionally, genes that encode calcium-transporting ATPases and other calcium pumps found on the plasma membrane were analyzed. In general, genes encoding calcium channels are most highly expressed in BSL cells relative to other sample groups, except for *ORAI2, SLC8A1, ATP2B1,* and *ATP2A2* (Figure 6A). *ORAI1* and *STIM1* show strong downregulation in Plasma Chase cells relative to cells from other experimental phases. Trajectories of gene expression of select calcium channel related genes (*ORAI1, ORAI2, ATP2B1, ATP2B4,* and *SLC8A1*) over pseudotime is represented in Figure 6B. This analysis indicates significant expression changes in select genes encoding calcium channels as a result of pump stimulation and interaction with SCD in a low iCa environment. *ATP2B1* has the most significant changes in gene expression through pseudotime, starting with an increase, followed by a decrease (Figure 5B). *SLC8A1*, which encodes a Sodium/Calcium Exchanger 1 (NCX1), starts with decrease in expression, followed by an increase. A heatmap of selected calcium-related GO pathways and enrichment of gene expression across the sample groups is shown in Figure 5C. In general, calcium-signaling and calcium channel related pathways are most upregulated in SCD bound cells relative to BSL and Circ cells. Consistent with the low iCa environment of SCD therapy, calcium ion binding is most enriched in BSL cells. Calcium channel activity is enriched significantly in elute cells relative to all other sample groups.

**Figure 6.**
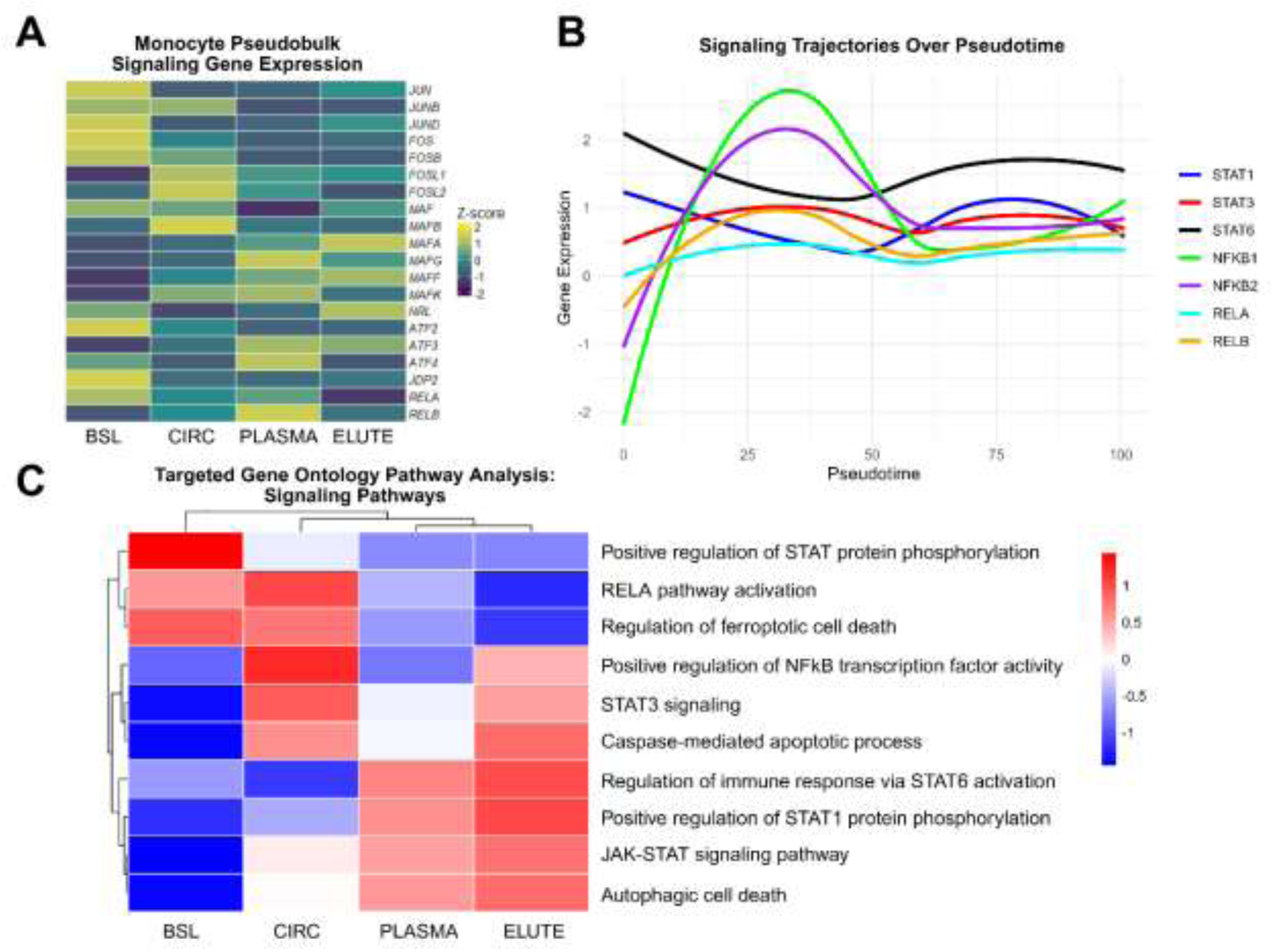
Dynamic changes in transcription factors AP-1 and NFkB related gene expression in monocytes by stage. **(A)** Heatmap depicting expression of targeted transcription factor related genes in pseudobulk of monocytes by stage. **(B)** Expression of individual transcription factor related genes plotted across pseudotime indicates *NFKB1* and *NFKB2* initially increase in unstimulated human blood at the onset of experimental exposure to SCD whereas *STAT1* and *STAT6* initially decrease. **(C)** Heatmap representing Gene Ontology: Biological Process pathways across BSL, Circ, Plasma, and Elute. Pathways were selected for their associations with AP-1 and NFkB related pathways.

### Results from Interrogation of Pathways Involving Critical Transcription Factors AP-1 and NFkB

Figure 7 analyzes gene expression of genes from the AP-1, NFkB, and STAT families across experimental phase. STAT, AP-1, and NFkB family proteins are critical for transcription of inflammatory genes (36–38). Gene expression of subunits of the AP-1 and NFkB transcription factors were interrogated across experimental phases in Figure 7A. AP-1 family genes were selected using a list by Bhosale PB et. al (36).

**Figure 7.**
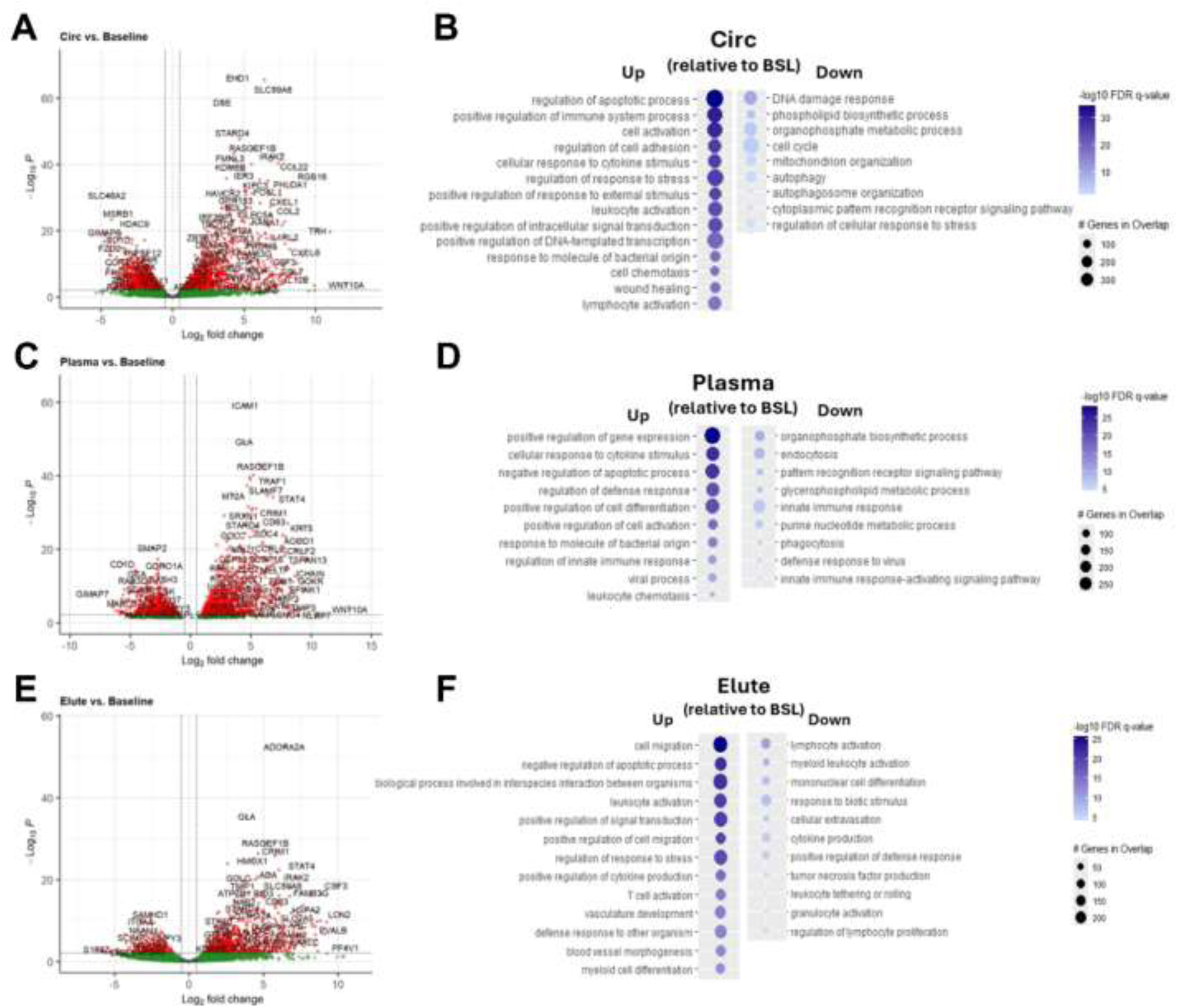
Differential impacts of individual experimental stages compared to Baseline (BSL). Volcano plots depicting gene expression comparing **(A)** Circ to BSL, **(C)** Plasma to BSL, and **(E)** Elute to BSL; threshold for significant genes, p < 0.01 and log2Fold > 0.5. Bubble plots representing up and down regulated (relative to BSL) GO: Biological Process pathways in **(B)** Circ, **(D)** Plasma, and **(F)** Elute stages. Pathways were manually selected to reduce redundancy and highlight pathways related to inflammation, infection, wound healing, cell cycle, metabolic reprogramming, autophagy, and/or apoptosis.

Expression is variable, with some genes (*JUN, JUNB, JUND, FOS,* and *FOSB)* most enriched in BSL cells, while other genes (*MAFB, MAFA, MAFG, MAFF,* and *MAFK)* were least enriched in BSL cells with variable expression patterns across the other sample groups. *MAF* was strongly downregulated in plasma chase cells relative to other experimental phases. *RELA* and *RELB,* subunits of the NFkB transcription factor complex, were also analyzed. *RELA* was most enriched in BSL cells and downregulated in Elute cells, while *RELB* was most enriched in plasma chase cells. *NFkB1, NFkB2, STAT1, STAT3,* and *STAT6* gene expression trajectories over pseudotime were analyzed in panel 7B. *NFkB1* and *NFkB2* show the most variable gene expression over the progression of pseudotime, starting with an increase followed by a decrease. *RELB* has a similar but less dramatic trajectory. *STAT1* and *STAT6* have similar trajectories, with a slight decrease followed by an increase. *RELA* expression appears to remain stable through the progression of pseudotime. These changes in expression over pseudotime suggest changes in transcription factor activity as a result of pump stimulation and SCD interaction. Panel 6E shows a heatmap of selected NFkB, STAT, and Cell Death GO pathways across experimental phase. Some cell death pathways, such as regulation of ferroptotic cell death and caspase-mediated apoptotic processes, are enriched in Circ cells relative to BSL, which could suggest that similar to previous findings regarding NE, promoting cell death pathways may also be involved in the immunomodulatory effects of SCD therapy on MO.

### Results from ToppGene Analysis of most differentially enriched pathways between experimental phases

ToppGene was used to analyze the most differentially expressed genes and gene pathways between experimental phases, starting with comparisons to BSL cells (Figure 8). Several cytokines/chemokines are depicted as being significantly enriched in Circ cells relative to BSL cells, including *CXCL1, CXCL6, CCL2, CCL7, CCL22, IL6,* and *IL12B* (Figure 8A). *FOSL1*, an AP-1 family gene, is also relatively enriched in Circ relative to BSL. Enrichment of top gene pathways as analyzed by ToppGene is represented in panel B. Circ cells, relative to BSL cells, had enrichment in both pro- and anti-inflammatory gene pathways, such as cell activation, positive regulation of immune system processes, LE activation, chemotaxis, and wound healing (Figure 8B). Circ cells also showed enrichment in apoptosis regulatory pathways. Upregulation of pro-inflammatory genes and gene pathways in Circ cells is an anticipated result with mechanical shear stress from the pump system, but the effects of SCD therapy and the low iCa environment may also be contributing to enrichment of certain genes and gene pathways due to release of bound cells during the reperfusion period.

**Figure 8.**
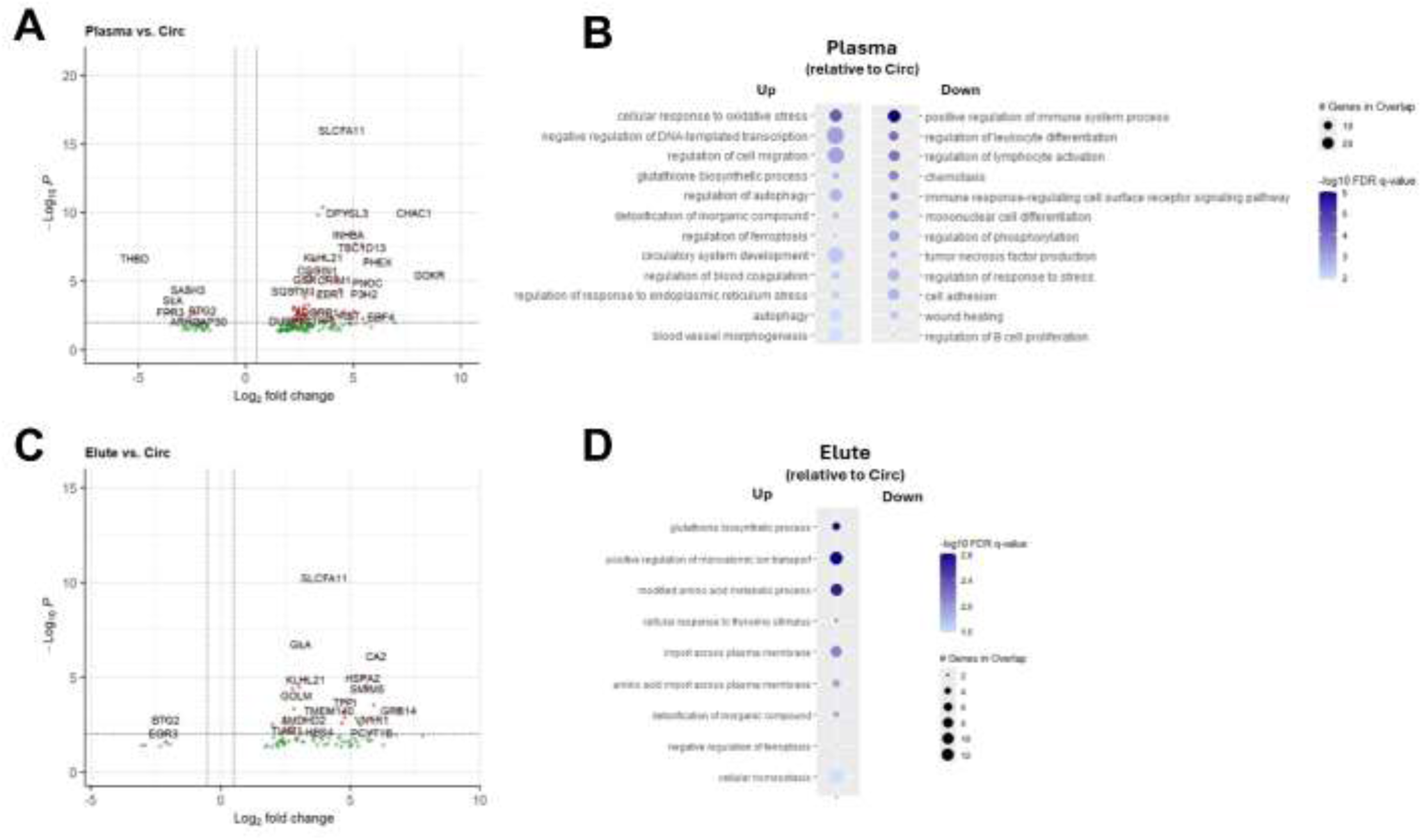
Differential impacts of individual experimental stages compared to Circ. Volcano plots depicting gene expression comparing **(A)** Plasma to Circ, and **(C)** Elute to Circ; threshold for significant genes, p < 0.01 and log2Fold > 0.5. Bubble plots representing up and down regulated (relative to BSL) GO: Biological Process pathways in **(B)** Plasma, and **(D)** Elute stages. Pathways were manually selected to reduce redundancy and highlight pathways related to inflammation, infection, wound healing, cell cycle, metabolic reprogramming, autophagy, and/or apoptosis.

Relative to Circ, BSL cells had enrichment in gene pathways involving cell cycle, metabolism, and autophagy. Panels 8C and 8D perform similar analysis as 8A and 8B, now examining enrichment in Plasma Chase cells relative to BSL cells. Pathways involving regulation of immune responses are enriched, including leukocyte chemotaxis, response to chemokine stimulus, regulation of cell activation, and regulation of immune responses (Figure 8D). BSL cells, relative to Plasma Chase cells, are enriched in endocytic pathways, such as endocytosis and phagocytosis. Panels 8E and 8F depict gene and gene pathways enriched in Elute cells relative to BSL cells. Elute cells show upregulation in gene pathways involving inflammation, including cell migration, LE activation, and positive regulation of cytokine production (Figure 8F).

In Figure 9, similar analysis using ToppGene was performed as in Figure 8, now interrogating gene expression relative to Circ samples. Plasma Chase cells showed enrichment in gene pathways involved in cellular response to stress, such as oxidative and endoplasmic reticulum stress, as well as regulatory pathways such as regulation of cell migration, autophagy, and ferroptosis (Figure 9B). Meanwhile, Circ cells were more enriched in immune response pathways relative to Plasma Chase cells, such as regulation of immune system process, cell adhesion, tumor necrosis factor (TNF) production, and chemotaxis (Figure 9B).

In cells Eluted from the SCD, metabolic and biosynthesis pathways were enriched, such as glutathione biosynthetic process and modified amino acid metabolic process (Figure 8D). Additionally, cells eluted from the SCD showed enrichment in transport, such as regulation of monoatomic ion transport and import across the plasma membrane.

## Discussion

SCD immunomodulatory activity is complex and multifaceted, making aspects of the MoA difficult to fully elucidate. The general concept is easy to follow as it is a fiber-filled, blood contacting device that can sequester adherent cell populations. The SCD appears to require a blood flow path along biocompatible membrane surfaces with low shear stress approximating capillary shear and in so doing, mimics a capillary bed where cells can be selectively addressed. The low ionized calcium (iCa) environment (<0.4mM) promoted with regional citrate anticoagulation is a key aspect of therapy which limits activation and reduces inflammation (13, 15). Under these conditions the SCD membrane was hypothesized to selectively bind the most activated neutrophils and monocytes due to the calcium dependency of binding reactions of cell surface integrins on the leukocyte.

SCD immunomodulatory therapy is currently directed toward treating acute indications like AKI, which are dominated by NE processes. However, in our experience to date, investigating chronic disease states, as well as tissue repair processes after acute injury, SCD may also play an important role via monocyte/macrophage immunomodulation. Understanding the SCD MoA for processing MO is integral for clinical translation into both acute and chronic tissue injury processes, including End Stage Renal Disease (ESRD). MO MoA revealed by scRNAseq and subsequent GSEA may demonstrate key gene programming shifts with SCD therapy in the MO cell population being returned to the patient’s circulation. More efficient SCD therapy may be developed with deeper understanding of these programming shifts as well identifying targets for pharmaceutical drug development.

As an initial foray into RNA sequencing, bulk RNAseq was used to compare basal (aka naïve) LE samples in normal and low iCa environments. In low iCa environments such as in the SCD with RCA, downregulation of the expression of several gene families was observed including the CCL subfamily (CCL2,7,8,12, 15 and 23) along with CCR7, the IL12 family cytokines(IL12, 27, 35), and CXCL9, CXCL10, CXCL11 (known as Mig, IP-10, and I-TAC) (data not shown). Single-cell RNA sequencing was utilized to further evaluate how SCD therapy alters the fate of MO after binding, sequestration, and release. The results of scRNAseq suggest that SCD binds the most inflammatory monocyte phenotypes. Supported by our previous data (19). scRNAseq shows that expression of genes encoding pro-inflammatory cytokines *TNF-α*, *IL6,* and *CXCL8* are most upregulated in cells sequestered by the SCD (Elute) relative to cells from other experimental phases (Figure 5A). Gene pathways involved in regulation of inflammation, such as ROS metabolic process, chemotaxis, and regulation of inflammatory response are more enriched in Elute cells relative to Circ cells (Figure 5B). Elute cells relative to BSL cells show enrichment in genes involved in cell migration, leukocyte activation, and cytokine production (Figure 8F). ScRNAseq data also shows that the cells released back into circulation are less inflammatory, shifting the proportion of monocyte subsets more towards those of anti-inflammatory phenotypes over time. SCD released cells in Plasma Chase and Circ samples show upregulation of the anti-inflammatory cytokine *IL-10* (Figure 5A). Interestingly, *IL-1RN,* the gene that encodes the IL-1 receptor antagonist (IL-1Ra) protein, was strongly enriched in SCD bound cells (Figure 5A). IL-1Ra can be induced in monocytes under pro-inflammatory conditions, such as activation via lipopolysaccharide (LPS) and adherence to immunoglobulin G (IgG) (39), so it is possible that the activation of monocytes in this model system may contribute to enrichment of *IL-1RN.* Additionally, recombinant IL-1Ra has shown promise as a therapeutic approach in treating pediatric patients with secondary hemophagocytic lymphohistocytosis (). Enrichment of *IL-1RN* in SCD sequestered cells may have a similar therapeutic effect, indicating this receptor antagonist as a potential key player in the immunomodulatory effects of SCD therapy.

Preferential binding of inflammatory MO to SCD and release of less inflammatory MO back into circulation is also supported by monocyte subset analysis. Unsupervised clustering of MO generated 11 clusters (Figure 2A). BSL cells appear to correlate most with clusters on the right side of the UMAP space, while Circ, Plasma Chase, and Elute cells appear to correlate most with clusters on the left (Figure 2B). Monocyte subsets are typically divided into pro-inflammatory classical and intermediate monocytes, and anti-inflammatory non-classical monocytes (3). Our analysis focused on classical and non-classical MO gene markers. BSL associated clusters 2, 3, 4, and 7 had the highest classical MO scores as determined by singleR, indicating a reduction in classical MO phenotype as a result of SCD therapy (Figure 2E) Cells in clusters 5 and 8 had low expression of the gene that encodes CD14 and increased expression of the gene that encodes CD16 (Figure 2D), had the highest non-classical MO scores, and had the lowest classical MO scores as determined by AddModuleScore (Figure 2E). Cluster 5 has significant downregulation in pro-inflammatory gene pathways (Figure 3C) as well as upregulation in some anti-inflammatory gene pathways (Figure 3D).

These are all phenotypic characteristics of non-classical monocytes. Additional evidence can be found when examining top gene markers of these clusters, which include *CDKN1C* and *LYPD2* for cluster 5, and *ADA, LLRC32*, and *CASP5* for cluster 8, genes that are enriched in non-classical MO relative to other immune cell types (40). Cluster 5 is interesting in its composition, with a relatively even mixture of BSL and Circ cells (Figure 4B). No other clusters have this even of a distribution of BSL/Circ cells. Cluster 8 is primarily composed of Circ cells, indicating additional cells with a non-classical MO phenotype in the Circ cell population (Figure 4B). This is suggestive that there is a population of cells at BSL with a non-classical MO phenotype, and that there is also a population of cells are modulated to or maintain a non-classical MO phenotype after pump activation and SCD therapy. Pseudotime analysis also provides support for this hypothesis. When initiated at BSL, pseudotime mirrors progression through the *in vitro* system, starting with BSL clusters on the right side of the UMAP, moving to Circ, Plasma Chase, and Elute associated clusters on the left side of the UMAP, finally ending at cluster 5 (Figure 4A). This progression may mirror the path of non-classical MO: a population of cells at BSL resembling non-classical monocytes moves through the *in vitro* system and interacts with SCD and returns back to a similar non-classical MO phenotype. In summary, scRNAseq data suggests that there is a population of cells that, after pump activation and SCD interaction, are modulated to or remain a non-classical MO phenotype.

Citrate anticoagulation in chronic dialysis patients when analyzed by scRNAseq and GSEA, has shown to alter several gene families in monocytes including those involved with cell cycle regulation, cellular metabolism and cytokine signaling (20) which may in part allow for the reprogramming of immune response at play in SCD therapy. Calcium-dependent pathways and channels show significant gene expression changes through the course of SCD therapy (Figure 6). The store-operated calcium entry (SOCE) mechanism resupplies store organelles (primarily the endoplasmic reticulum) with Ca2+ after they have been depleted The influx of calcium occurs primarily through calcium-release activated calcium (CRAC) channels, which are composed of ORAI and STIM family members (20). Gene expression levels of ORAI and STIM are influenced by cell type, activation state, and in monocytes, reactive oxygen species (ROS) production (20, 21). Also involved in calcium homeostasis are plasma membrane Ca2+ ATPases (PMCA) that couple ATP hydrolysis with cellular efflux of Ca2+ ions (22), mitochondrial calcium uptake channels (41), and Na+/Ca2+ exchanger 1 (NCX1) (23). NCX1 can act in forward or reverse directions; under normal physiological conditions it acts in the forward direction, resulting in an influx of Na+ ions and an efflux of Ca2+ ions (23). When human monocytes/macrophages are incubated in Na+-free medium, NCX1 acts in the reverse direction which resulted in a release of pro-inflammatory cytokine TNF-α, suggesting NCX1 activity is influenced by extracellular conditions and may be associated with an inflammatory response in these cells (23). In general, expression of genes encoding calcium channels was most enriched in BSL cells, (Figure 6A). consistent with other experimental phases being exposed to a low iCa environment as a result of RCA. This also is supported by gene pathway analysis, with calcium ion binding being significantly upregulated in BSL cells relative to other experimental phases (Figure 6C). *SCL8A1,* which encodes the sodium/calcium exchanger NCX1, did not follow this pattern and was most enriched in Circ cells (Figure 6A). This may indicate increased efflux of intracellular Ca^2+^ ions in SCD treated MO and reduced inflammatory response as a result of the maintained low iCa environment intra- and extra-cellularly (42).

AP-1, NFkB, and STAT family proteins are heavily involved in transcriptional regulation and signal transduction of various inflammatory genes and gene pathways (43).Signaling pathways involving NFkB and AP-1 control various cellular processes, including inflammation and apoptosis (36). The AP-1 transcription factor complex is formed by homo- and heterodimerization of JUN, FOS, or ATF subunits (44). The AP-1 transcription factor can be activated in response to pro-inflammatory cytokines and depending on the composition of its subunits, can target a variety of genes and cell processes (45). The FOS subunit of the AP-1 transcription factor was strongly upregulated in clusters 6 (Figure 2C). Additional AP-1 family members show significant expression changes across experimental groups, indicating AP-1’s involvement in differential expression of genes between these groups. Previous findings by our group suggest that SCD therapy promotes cell-death in neutrophils (19), so analysis of AP-1, NFkB, and STAT family proteins was preformed to analyze the activity of these regulatory genes and of cell-death pathways in MO. Ferroptosis is an autophagy-dependent form of programmed cell death (46). Activation of STAT and KFkB family proteins, many members of which are enriched in SCD treated cells, promote ferroptosis (46). Circ cells show enrichment in ferroptosis related pathways relative to other experimental phases (Figure 7C). Additionally, Elute cells relative to Circ cells showed upregulation of genes involved in synthesis of glutathione, a ferroptosis inhibitor (46). Ferroptosis, therefore, may be another immunomodulatory effect of SCD therapy that contributes to reduced proportion of inflammatory MO in circulation.

The data presented has several limitations. Gene expression changes seen in Circ samples relative to BSL samples could be attributed to stimulation via the blood pump, or SCD therapy. However, the lack of a control sample without SCD treatment makes it difficult to attribute differential expression to SCD therapy alone. Additionally, Plasma Chase and Elute samples had significantly lower cell numbers relative to BSL and Circ samples, making analysis of gene expression changes in these groups much less robust. Identification of monocytes by singleR also shows potential intermixing of other cell types (such as neutrophils) due to overlapping gene expression profiles (Supplemental Figure 1). Donor blood used for the *in vitro* experiment was from healthy donors. Hyperinflammatory disease states may alter phenotypic/genotypic profiles of LE, and pump stimulation of LE is not a perfect substitute for mimicking these disease states. Additionally, post-transcriptional and post-translational regulation of gene and protein expression are not accounted for in scRNAseq analysis. Despite these limitations, the data presented provides strong evidence that SCD preferentially binds inflammatory (classical) monocytes and releases and/or modulates MO to a less inflammatory non-classical phenotype in combination with the low iCa environment generated by RCA. There is also evidence that SCD therapy may promote programmed cell death via ferroptosis in MO, similar to the effects of SCD therapy on NE. Future studies should aim to analyze clinical samples, with a focus on systemic changes in MO phenotypes.

## Conclusions

Utilizing the scRNAseq as a tool to interrogate the mechanism of SCD’s cell directed therapy, has both confirmed previous mechanistic details gleaned from cell surface and functional testing, bolstering understanding of autologous leukocyte processing, but it also has informed new avenues of inquiry. This work points to some specific regulatory genes such as Fos, which may be an important target of SCD therapy.

Further investigating nuances of the cell-directed approach by scRNAseq may help elucidate why SCD has a beneficial clinical effect to immunomodulate, but not immunosuppress, complex dysregulated inflammatory processes.

## Acknowledgements

This work was supported in part by funding from NIH NCATS 2R44 TR001324. We would like to thank Dr Liandi Lou who helped conduct aspects of the IVBC studies and preparation of cell samples ahead of scRNAseq. We would also like to thank the University of Michigan (UM) Biomedical Research Core Facilities (BRCF) Advanced Genomics Core (AGC) for their aid in running scRNAseq. The Applied Systems Biology Core (ASBC) and Data Analytics Services Core (DASC), specifically Wenjun Ju, Celine Berthier and Edgar Otto in early consultation on the set up and conduct of these studies, as well as Jeff Beamish and Jeff Hodkin.

## Author Contributions

Conceptualization: CJP, HDH, JO

Data curation: CJP, JO

Formal Analysis: CJP, JO

Funding acquisition: CJP

Investigation: CJP, HDH, JO

Methodology: CJP, JO

Project administration: HDH, CJP

Resources: HDH, CJP

Supervision: HDH, CJP

Validation: CJP, HDH, JO

Writing: CJP, HDH, JO

Writing-review & editing: CJP, HDH, JO

## Data Availability Statement

As much as possible, data has been made available directly within the text, figures and legends of the manuscript. If there is additional data required during the review process, the authors plan to submit additional supplemental materials to address those requests.

## Additional Information (Competing Interests Statement)

HDH and CJP are shareholders of SeaStar Medical, a publicly traded company commercializing SCD technology. HDH, CJP and TM are employees of Innovative Biotherapies.

