## Supplemental Materials for "Immunomodulatory Effects of a Cell Processing Device: Insights from Single Cell RNA Sequencing and Gene Set Enrichment Analysis"

**Supplementary File 1**

| SingleR Cell Type | Number of Cells |
| --- | --- |
| B_cell | 310 |
| CMP | 215 |
| DC | 58 |
| Epithelial_cells | 3 |
| GMP | 20 |
| HSC_-G-CSF | 160 |
| HSC_C34+ | 13 |
| Keratinocytes | 2 |
| Macrophage | 450 |
| MEP | 48 |
| Monocyte | 25877 |
| Myelocyte | 9 |
| Neutrophils | 5723 |
| NK_cell | 1221 |
| Platelets | 865 |
| Pre-B_cell_CD34- | 100 |
| Pro-B_cell_CD34+ | 1 |
| T_cells | 2894 |

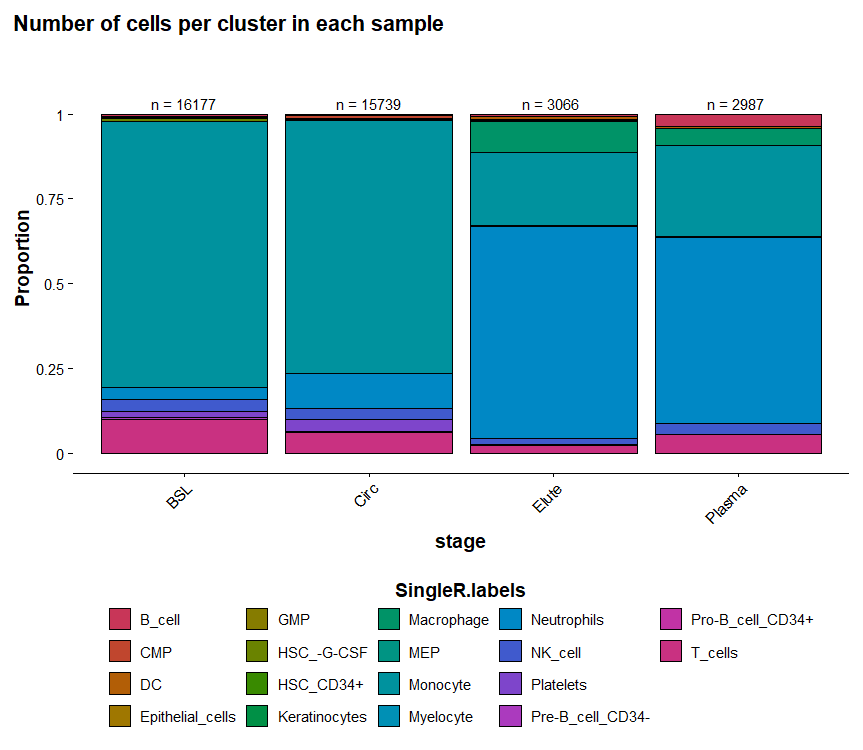

**Supplemental Figure 1: Cell type assignments by SingleR based on stage of *in vitro* experiment.**

**Figure 3F GO terms**

| GO Term | Description |
| --- | --- |
| GO:0050729 | Positive Regulation of inflammatory response |
| GO:0045087 | Innate immune response |
| GO:0072593 | Reactive oxygen species metabolic process |
| GO:0045059 | Positive regulation of T cell activation |
| GO:0019882 | Antigen processing and presentation |
| GO:0032755 | Positive regulation of interleukin-6 production |
| GO:0071222 | Cellular response to lipopolysaccharide (LPS) |
| GO:0006935 | Chemotaxis |
| GO:0034599 | Cellular response to oxidative stress |
| GO:0034142 | Toll-like receptor signaling pathway |

**Figure 3G GO terms**

| GO Term | Description |
| --- | --- |
| GO:0050727 | Regulation of inflammatory response |
| GO:0010594 | Positive regulation of cell-matrix adhesion |
| GO:0030889 | Negative regulation of B cell activation |
| GO:0032723 | Positive regulation of interleukin-10 production |
| GO:0006909 | Phagocytosis |
| GO:0002244 | Hemostasis |
| GO:0032740 | Positive regulation of interleukin-4 production |
| GO:0030198 | Extracellular matrix organization |
| GO:0042060 | Wound healing |
| GO:0001525 | Angiogenesis |

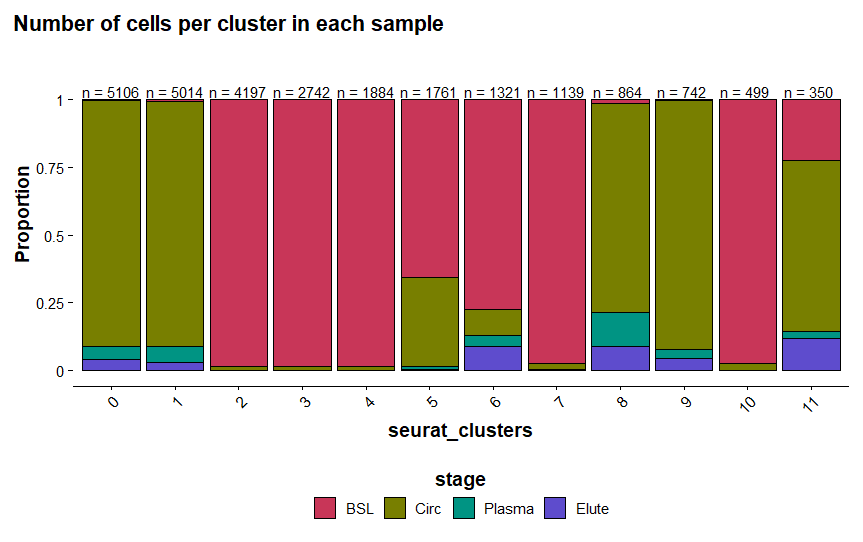

**Supplemental Figure 2: Breakdown of monocyte Clusters by stage of *in vitro* experiment**

**Figure 4B GO Terms**

| GO Term | Description |
| --- | --- |
| GO:0045059 | Positive regulation of T cell activation |
| GO:0019882 | Antigen processing and presentation |
| GO:0050727 | Regulation of inflammatory response |
| GO:0006909 | Phagocytosis |
| GO:0032723 | Positive regulation of interleukin-10 production |
| GO:0030889 | Negative regulation of B cell activation |
| GO:0045087 | Innate immune response |
| GO:0010594 | Positive regulation of cell-matrix adhesion |
| GO:0050729 | Positive regulation of inflammatory response |
| GO:0032755 | Positive regulation of interleukin-6 production |
| GO:0034142 | Toll-like receptor signaling pathway |
| GO:0072593 | Reactive oxygen species metabolic process |
| GO:0002244 | Hemostasis |
| GO:0034599 | Cellular response to oxidative stress |
| GO:0032740 | Positive regulation of interleukin-4 production |
| GO:0030198 | Extracellular matrix organization |
| GO:0071222 | Cellular response to lipopolysaccharide (LPS) |
| GO:0006935 | Chemotaxis |
| GO:0042060 | Wound healing |
| GO:0001525 | Angiogenesis |

**Figure 5C GO Terms**

| GO Term | Description |
| --- | --- |
| GO:0005509 | Calcium ion binding |
| GO:0046777 | Calcium-dependent protein autophosphorylation |
| GO:0004698 | Calcium-dependent kinase activity |
| GO:0070588 | Calcium ion transmembrane transport |
| GO:0005262 | Calcium channel activity |
| GO:0050848 | Regulation of calcium-mediated signaling |
| GO:0071297 | Calcium-mediated signaling involved in inflammatory response |
| GO:0030003 | Regulation of calcium ion transport |
| GO:0006874 | Calcium ion homeostasis |
| GO:0034220 | Calcium-dependent activation of MAPK activity |

**Figure 6C GO Terms**

| GO Term | Description |
| --- | --- |
| GO:0038128 | Positive regulation of STAT protein phosphorylation |
| GO:0038089 | RELA pathway activation |
| GO:1901214 | Regulation of ferroptotic cell death |
| GO:0051092 | Positive regulation of NFkB transcription factor activity |
| GO:0038001 | STAT3 signaling |
| GO:004381 | Caspase-mediated apoptotic process |
| GO:0043953 | Regulation of immune response via STAT6 activation |
| GO:000038126 | Positive regulation of STAT1 protein phosphorylation |
| GO:0007259 | JAK-STAT signaling pathway |
| GO:0010506 | Autophagic cell death |
